# Protein Disulfide Isomerases Control the Secretion of Wnt proteins

**DOI:** 10.1101/429845

**Authors:** Nanna Torpe, Sandeep Gopal, Oguzhan Baltaci, Lorenzo Rella, Ava Handley, Hendrik C. Korswagen, Roger Pocock

## Abstract

Appropriate Wnt morphogen secretion is required to control animal development and homeostasis. Although correct Wnt globular structure is essential for secretion, proteins that directly mediate Wnt folding and maturation are incompletely characterized. Here, we report that protein disulfide isomerase-1 (PDI-1), a protein folding catalyst and chaperone, controls secretion of the *Caenorhabditis elegans* Wnt ortholog EGL-20. We find that PDI-1 function is required to correctly form an anteroposterior EGL-20/Wnt gradient during embryonic development. Further, PDI-1 performs this role in EGL-20/Wnt-producing epidermal cells to cell-non-autonomously control EGL-20/Wnt-dependent neuronal migration. Using pharmacological inhibition, we further show that PDI function is required in human cells for Wnt3a secretion, revealing a conserved role for disulfide isomerases. Together, these results demonstrate a critical role for PDIs within Wnt-producing cells to control long-range developmental events that are dependent on Wnt secretion.

## Introduction

Wnt glycoproteins are predominant regulators of metazoan cell fate patterning and tissue assembly. During development, Wnt proteins can form long-range concentration gradients that provide positional and instructive information to surrounding cells and tissue (Cadigan et al., 1998; Clevers, 2006). The scale and amplitude of Wnt protein gradients requires tight regulation, as defects in gradient formation are linked to developmental deficits and disease (Christodoulides et al., 2006; MacDonald et al., 2014; Person et al., 2010; Roifman et al., 2015). As such, there is substantial interest in understanding how Wnt proteins are synthesized, modified, secreted and delivered to receiving cells.

Wnt proteins are characterized by 24 invariantly positioned cysteine residues that form intramolecular disulfide bonds to maintain functional Wnt globular secondary structure (Miller, 2002; Willert and Nusse, 2012). Synthesis and maturation of Wnt proteins occur in the endoplasmic reticulum (ER) and Golgi apparatus, facilitated by numerous processing enzymes and molecular chaperones (Braakman and Bulleid, 2011; Willert and Nusse, 2012). During Wnt protein maturation, complex folding events and posttranslational modifications occur that are required for secretion and formation of a Wnt concentration gradient (Janda et al., 2012; Willert and Nusse, 2012). Biochemical characterization of secreted Wnt revealed that Wnt proteins undergo functionally important posttranslational modifications such as lipidation and glycosylation (Caramelo and Parodi, 2007; Takada et al., 2006; Willert et al., 2003). A particularly critical step for Wnt protein maturation and secretion is acylation of a specific serine residue by the membrane-bound O-acyltransferase called Porcupine (Kadowaki et al., 1996; Vandenheuvel et al., 1993). Inhibiting Wnt protein acylation ablates Wnt signaling and results in severe developmental defects in animal models and causes human disorders such as focal dermal hypoplasia (Barrott et al., 2011; Biechele et al., 2011; Grzeschik et al., 2007; Wang et al., 2007). Transfer of the acyl group to Wnt proteins is required for a functional interaction with the seven-pass transmembrane protein Wntless (Wls), which binds to and conveys Wnt proteins to the cell surface of Wnt-producing cells. The transport of Wnt proteins to the cell surface by Wntless is critical for correct Wnt protein secretion (Banziger et al., 2006). Following delivery of Wnt protein, Wls is recycled to the Golgi via endosomes and the retromer complex where it is able to escort a newly synthesized Wnt protein to the cell surface (Belenkaya et al., 2008; Coudreuse et al., 2006; Port et al., 2008; Prasad and Clark, 2006; Yang et al., 2008). In *Caenorhabditis elegans*, studies have also shown that MIG-14/Wls and retromer complex components such as VPS-35 are required for appropriate formation of an EGL-20/Wnt gradient (Coudreuse et al., 2006; Yang et al., 2008). Further, in the absence of retromer complex function, MIG-14/Wls is degraded in lysosomes, thereby limiting EGL-20/Wnt secretion (Yang et al., 2008). Therefore, key molecular components and mechanisms that control Wnt secretion are highly conserved across metazoa.

Wnt proteins are known to play major roles in directing neuronal migration and axon-dendritic guidance in vertebrates (Bocchi et al., 2017; Yoshikawa et al., 2003). In *C. elegans* Wnt proteins have also been shown to control specific neurodevelopmental events, including neuron migration, neuronal polarity and axon guidance (Coudreuse et al., 2006; Hilliard and Bargmann, 2006; Mentink et al., 2014; Modzelewska et al., 2013; Pan et al., 2006). Further, control of EGL-20/Wnt gradient formation by MIG-14/Wls and the retromer complex is required to drive development of the nervous system (Coudreuse et al., 2006; Yang et al., 2008). We therefore employed the correlation between correct Wnt gradient formation and neuronal development in the *C. elegans* model to identify novel regulators of Wnt secretion.

Wnt proteins are rich in disulfide bonds between cysteine side chains that are mostly formed during translation in the ER (Fass, 2012). These disulfide bonds enhance Wnt stability in the proteolytic and oxidizing extracellular environment to enable appropriate cell to cell communication. Therefore, the coordination of disulfide bonds is a vital and pervasive step during Wnt biogenesis (MacDonald et al., 2014). Previous studies demonstrated that erroneous disulfide bonds can form during Wnt synthesis and that correct disulfide bond formation is required for Wnt secretion (MacDonald et al., 2014; Zhang et al., 2012). This suggests that regulatory enzymes that monitor and shuffle disulfide bonds would be essential for appropriate Wnt secretion and signaling. However, until now, the identity of these regulatory molecules has not been revealed. Here, we identify protein disulfide isomerases (PDIs) as master controllers of Wnt secretion. Our genetic and pharmacological characterization of *C. elegans* PDI-1 reveals that it controls EGL-20/Wnt-directed neuronal migration from EGL-20-producing epidermal cells. In the *C. elegans* embryo, we find that PDI-1 is required for EGL-20-Wnt gradient formation in the temporal window that neuronal migration occurs. Finally, we reveal that PDI function is conserved, as pharmacological inhibition of PDI in human cells specifically impairs Wnt secretion. Taken together, we have identified PDIs as novel and pervasive regulators of Wnt secretion. This provides an added level of complexity to the control of Wnt secretion and posits PDIs as potential targets for therapeutic regulation of Wnt-dependent cell signaling.

## Results

### PDI-1, a Protein Disulfide Isomerase, Regulates Neuron Migration

EGL-20/Wnt is expressed and secreted from a discrete subset of hypodermal (epidermal) and muscle cells in the posterior of *Caenorhabditis elegans* (Whangbo and Kenyon, 1999) (Figure 1A). EGL-20 function is critical for specific neurodevelopmental events, including anterior migration of the hermaphrodite-specific neurons (HSNs) (Desai et al., 1988) (Figure 1A). We therefore used HSN development as a readout of EGL-20 function to identify molecules that control Wnt maturation and secretion (Figure 1). Protein disulfide isomerases (PDIs) are a family of protein chaperones that reside in the ER to catalyze the formation (oxidation), breakage (reduction), and rearrangement (isomerization) of disulfide bonds (Wilkinson and Gilbert, 2004). Due to the importance of disulfide bond formation for Wnt secretion and function (MacDonald et al., 2014; Zhang et al., 2012), we hypothesized that PDIs may be important for EGL-20-directed HSN development. Using RNA-mediated interference (RNAi), we knocked down the expression of the five *C. elegans* PDI-encoding genes (*pdi-1, pdi-2, pdi-3, pdi-6* and *C14B9.2*) and analyzed HSN development (Figure 1B-C). This screen identified a specific requirement for *pdi-1* during HSN development, where ∼20% of *pdi-1(RNAi)*-treated animals exhibit HSN undermigration defects (Figure 1B-C). We confirmed the role of *pdi-1* in HSN development using a previously isolated *pdi-1(gk271)* null allele (Figures 1D-E, 2A, S1 and Table S1). As PDI proteins act as folding catalysts and molecular chaperones, we hypothesized that elevated temperature would exacerbate improper folding events and adversely affect HSN development (Mogk et al., 2002). We therefore examined *pdi-1(gk271)* mutant animals exposed to temperature stress and found that HSN developmental defects progressively increase with elevated incubation temperature, with animals incubated at 25°C exhibiting a penetrance of ∼50% (Figure S1). EGL-20 controls HSN migration during embryogenesis. To determine whether PDI-1 can also control EGL-20-dependent postembryonic neuronal development we examined the post-embryonically migrating QR.pax cell (Harris et al., 1996a). We found that indeed *pdi-1(gk271)* mutant animals exhibit defective migration of QR.pax (Figure S2). Taken together, we show that PDI-1 controls two temporally-distinct EGL-20-regulated migration events.

**Figure 1.**
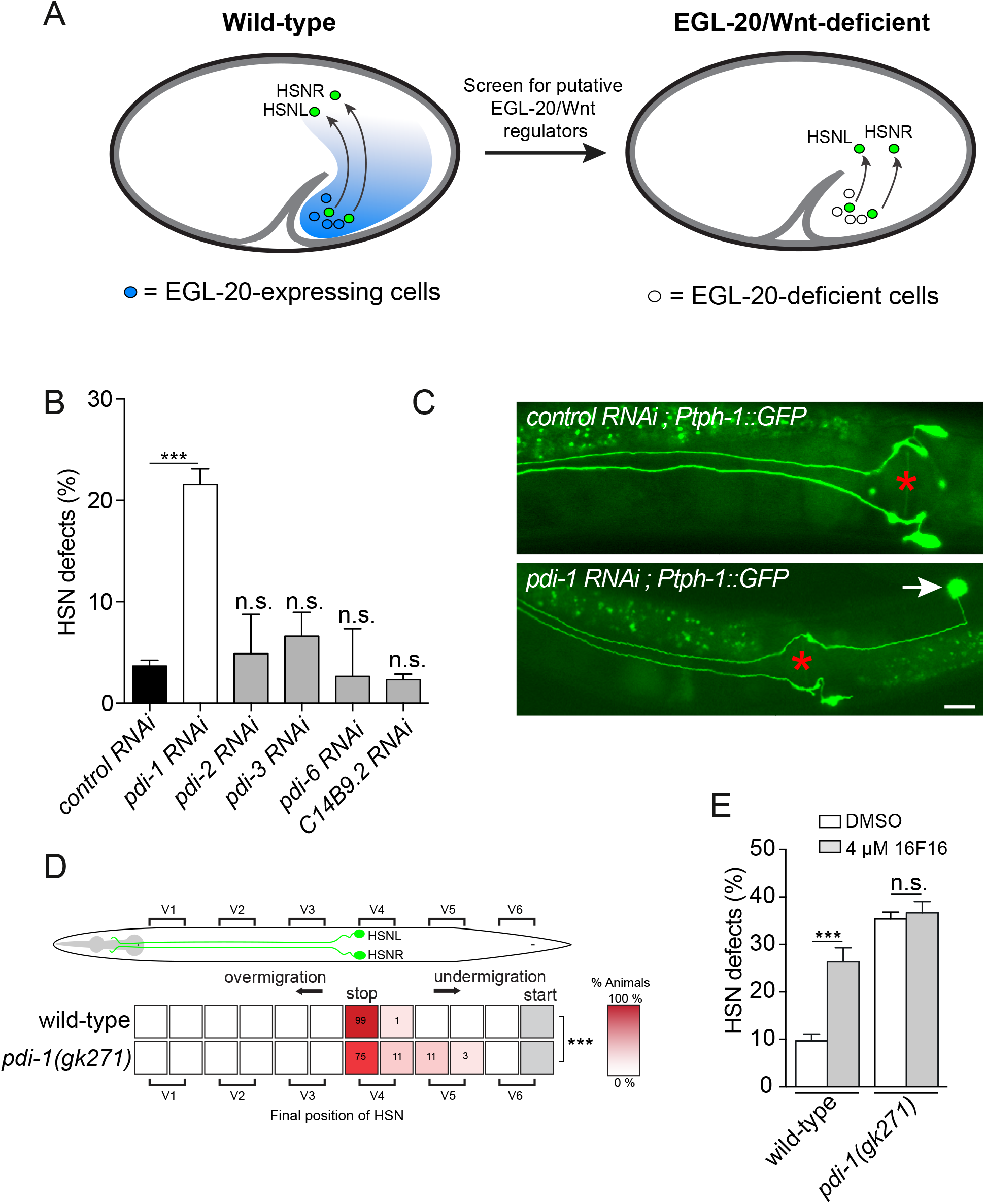
PDI-1, a protein disulfide isomerase, controls HSN developmen. (A) EGL-20/Wnt is expressed in a discrete number of cells in the posterior of the *C. elegans* embryo (blue circles, left embryo) and forms a long-range morphogen gradient (blue shading). Formation of the EGL-20/Wnt gradient is required for correct migration of the hermaphrodite-specific neurons (HSNL and HSNR) during embryogenesis from the posterior to the mid-body (green circles). Embryos with reduced gradient or complete loss of EGL-20/Wnt exhibit HSN migration defects (right embryo), which provides a screening platform for novel EGL-20/Wnt regulatory molecules. (B) Reduced function of *pdi-1*, but no other *pdi* genes, causes HSN developmental defects. HSN development was scored using a *Ptph-1::gfp* transgene (*zdIs13*). Data are expressed as mean ± s.d. and statistical significance was assessed by ANOVA followed by Dunnett’s Multiple Comparison Test by comparison to vector control RNAi. n>50, ***p<0.001, n.s. – not significant. (C) HSN anatomy of control RNAi (upper image) and *pdi-1* RNAi (lower image) young adult animals. HSN cell bodies migrate to a position posterior to the vulva (marked with red asterisk) and extend axons in a highly stereotypical manner. HSN axons extend ventrally and then around the vulva before entering the ventral nerve cord, where each axon is separated by the hypodermal ridge. The lower image is the most common phenotype observed when *pdi-1* expression is abrogated (see Table S1 for full details), where an HSN cell body undermigrates (white arrow). Ventral view, anterior to the left. Scale bar 20 μm. (D) Average position of the HSNs with respect to seam cells V1.a to V6.p in wild-type and *pdi-1(gk271)* animals (scored by Nomarski optics). Values listed are percentiles of the total number of cells scored; the red coded heat map displays the range of percentile values. Statistical significance was calculated using Fisher’s exact test. n>30 for each cell-type scored, *** p<0.0001. (E) Quantification of HSN developmental defects after exposure of wild-type and *pdi-1(gk271)* animals to the PDI small-molecule inhibitor, 16F16 (4 µM). Data are expressed as mean ± s.d. and statistical significance was assessed by ANOVA followed by Dunnett’s Multiple Comparison Test. n>50, ***p<0.001, n.s. – not significant.

*In vitro* studies previously identified small-molecule inhibitors of PDI proteins in mammalian cells (Hoffstrom et al., 2010; Kaplan et al., 2015). Using the PDI-specific 16F16 inhibitor, we asked whether manipulation of PDI activity could affect PDI-regulated HSN development *in vivo*. We incubated late larval stage 4 (L4) hermaphrodites in 4µM 16F16 for 48 hours and examined HSN development in the resultant progeny. We found that 16F16 causes HSN defects in wild-type animals similar to that caused by the *pdi-1(gk271)* mutation (Figure 1E). In contrast, when *pdi-1(gk271)* mutants were exposed to the PDI inhibitor we observed no enhancement of the HSN phenotype (Figure 1E). This suggests that 16F16 causes HSN developmental defects through inhibition of PDI-1 activity *in vivo*.

### PDI-1 Acts Cell-Non-autonomously From EGL-20/Wnt-expressing Epidermal Cells to Control Neuronal Migration

HSN migration occurs during mid-embryogenesis, between the comma and 1.5 fold stages (Desai et al., 1988). To determine the spatiotemporal expression pattern of *pdi-1* during embryogenesis, we constructed a transgenic reporter in which an 841 bp sequence upstream of the *pdi-1* ATG (up to the 5’ end of the preceding gene) drives expression of green fluorescent protein (GFP) (Figure 2A-I). We first detected GFP expression at the comma stage in hypodermal cells and expression in the hypodermis continues throughout embryogenesis (Figure 2B-I). The absence of detectable neuronal expression suggested that *pdi-1* acts cell-non-autonomously to regulate HSN development, which is a feature of other guidance pathways in *C. elegans* (Kennedy et al., 2013; Pedersen et al., 2013). To investigate the PDI-1 focus-of-action, we first drove *pdi-1* cDNA under the endogenous *pdi-1* promoter in *pdi-1(gk271)* mutant animals and analyzed HSN development (Figure 2J). As expected, expressing *pdi-1* under the control of the *pdi-1* promoter fully rescued the *pdi-1(gk271)* HSN developmental defects (Figure 2J). To determine if hypodermal expression of *pdi-1* is sufficient to rescue these defects we used the hypodermal-specific *dpy-7* promoter and found that the HSN defects of *pdi-1(gk271)* mutant animals were also fully rescued (Figure 2J). PDI proteins act in the ER lumen to catalyze protein folding and inhibit misfolded protein aggregation (Wilkinson and Gilbert, 2004). PDI proteins, including PDI-1, contain a C-terminal ER-retention signal (His-Glu-Glu-Leu or HEEL) that helps maintain their functional locale. We therefore examined whether the HEEL motif is required for PDI-1 regulation of HSN development. We found that deleting the ER retention signal (-HEEL) abrogates the ability of PDI-1 to rescue the HSN developmental defects of *pdi-1(gk271)* mutant animals (Figure 2J). Together, our data show that PDI-1 acts within the ER of hypodermal cells to cell-non-autonomously regulate HSN development.

**Figure 2.**
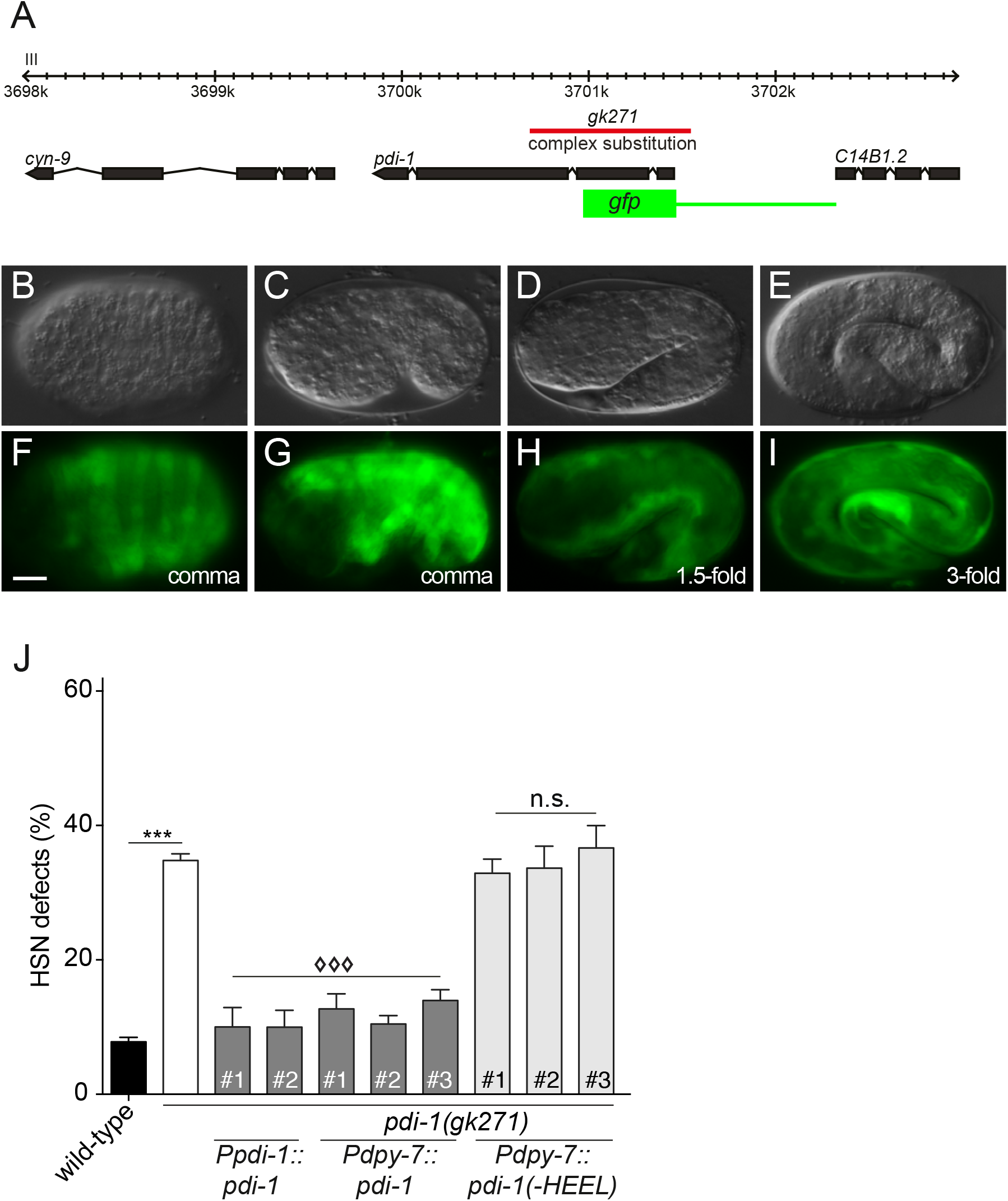
PDI-1 regulates HSN development non-cell-autonomously from the hypodermis. (A) Genetic locus of *pdi-1* showing gene structure in black and *gk271* genetic lesion in red. The 841 bp promoter used to drive *gfp* for expression analysis is in green. (B-I) Expression of a *Ppdi-1::gfp* transcriptional reporter during embryonic development. Expression is first observed at the comma stage (F – dorsal, G – lateral) in the hypodermis and continues to be expressed in this tissue in 1.5-fold and 3-fold embryos (H-I). Upper panels (B-E) are Nomarski micrographs and bottom panels (FI) are fluorescence images of the same embryos. Anterior to the left. Scale bar 10 μm. (J) HSN developmental defects of *pdi-1(gk271)* mutant animals are rescued by transgenic expression of *pdi-1* cDNA controlled by its own promoter and the hypodermal-specific *dpy-7* promoter. Deletion of the ER retention motif (-HEEL) abrogates PDI-1 rescuing ability in the hypodermis. Data are expressed as mean ± s.d. and statistical significance was assessed by ANOVA followed by Dunnett’s Multiple Comparison Test. n>50, ***p = 0.001 for wild-type compared to *pdi-1(gk271)*, ◊◊◊p<0.001 and n.s. – not significant for *pdi-1(gk271)* compared to rescue lines. # refers to independent transgenic lines. Scale bar 10 μm.

The HSN migration defects of *pdi-1(gk271)* mutant animals are comparable, albeit less severe, to phenotypes observed in *egl-20*/Wnt loss-of-function animals (Figure 3A, B, F and Table S1) (Forrester et al., 2004). In addition, *pdi-1* and *egl-20* are co-expressed in hypodermal tissue, with *egl-20* expression in a more restricted posterior region (Prasad and Clark, 2006; Whangbo and Kenyon, 1999). If PDI-1 functions to control EGL-20-directed HSN development, one may expect *pdi-1* to act in *egl-20*-producing hypodermal cells. We confirmed that the previously published 1900 bp *egl-20* promoter drives expression in a discrete subset of cells in the tail of embryos that temporally coincides with HSN migration (Figure 3C-D) (Harris et al., 1996b; Sulston et al., 1983). We used this promoter to drive *pdi-1* cDNA in *egl-20*-producing cells and found that the HSN developmental defects of *pdi-1(gk271)* mutant animals were fully rescued (Figure 3E). In contrast, expressing *pdi-1* cDNA using the promoter of *lin-44*, a Wnt-encoding gene expressed in an adjacent and non-overlapping subset of cells in the *C. elegans* posterior (Coudreuse et al., 2006; Klassen and Shen, 2007), was unable to rescue the HSN defects (Figure 3E). Thus, PDI-1 acts specifically from EGL-20-producing hypodermal cells to control HSN development.

**Figure 3.**
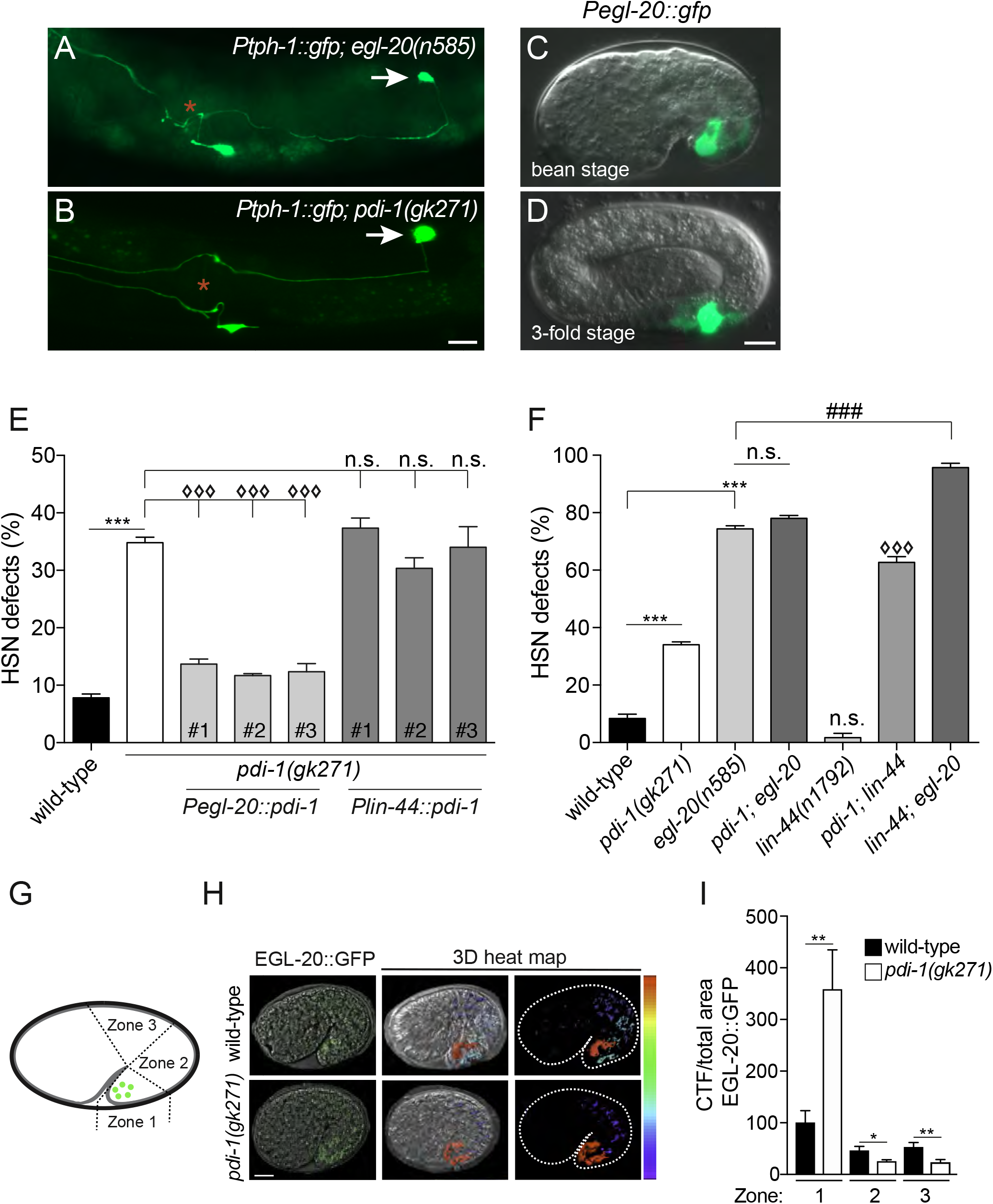
PDI-1 acts in the EGL-20/Wnt pathway to direct HSN development. (A-B) HSN anatomy of *egl-20(n585)* and *pdi-1(gk271)* mutant animals, where one of the HSN cell bodies undermigrates (white arrows). Vulval position is marked with a red asterisk. Ventral view, anterior to the left. Scale bar 20 μm. (C-D) Expression of a *Pegl-20::gfp* transcriptional reporter at the comma and 3-fold stages of embryonic development. As previously reported the *egl-20* reporter drives expression in a discrete subset of hypodermal and muscle cells in the posterior (Prasad and Clark, 2006; Whangbo and Kenyon, 1999). Anterior to the left. Scale bar 10 μm. (E) HSN developmental defects of *pdi-1(gk271)* mutant animals are rescued by transgenic expression of *pdi-1* cDNA controlled by the *egl-20* promoter (used in C-D) but not by the *lin-44* promoter, which expresses more posteriorly (Klassen and Shen, 2007). Data are expressed as mean ± s.d. and statistical significance was assessed by ANOVA followed by Dunnett’s Multiple Comparison Test. n>50, ***p = 0.001 for wild-type compared to *pdi-1(gk271)*, ◊◊◊p<0.001 and n.s. – not significant for *pdi-1(gk271)* compared to rescue lines. # refers to independent transgenic lines. (F) Double mutant analysis of Wnt morphogen mutant strains in combination with *pdi-1(gk271)*. The penetrance of *pdi-1(gk271); egl-20(n585)* mutant animals is not significantly different from the *egl-20(n585)* single mutant. Whereas, *lin-44(n1792)* double mutant combinations with either *pdi-1(gk271)* or *egl-20(n585)* exhibit an additive effect. See Table S1 for detailed phenotypic scoring. Data are expressed as mean ± s.d. and statistical significance was assessed by ANOVA followed by Dunnett’s Multiple Comparison Test. n>50, ***p<0.001 for wild-type compared to *pdi-1* and *egl-20* mutants, ◊◊◊p<0.001 for *pdi-1(gk271)* compared to *pdi-1(gk271); lin-44(n1792)* double mutant, ###p<0.001 for *egl-20(n585)* compared to the *egl-20(n585); lin-44(n1792)* double mutant and n.s. – not significant. (G-I) EGL-20::GFP expression and secretion analysis. (G) Schematic of a comma stage embryo expressing EGL-20::GFP from a group of posterior cells (green circles). Zones 1–3 represent the areas considered for measuring EGL-20::GFP levels using a corrected total fluorescence calculation (see Experimental Procedures). (H) 3D reconstruction and heat map of wild-type and *pdi-1(gk271)* mutant comma stage embryos expressing EGL-20::GFP. Single slice of a Z-stack from confocal microscopy (left) and 3D reconstructed heat maps of the of the same embryos (center and right). EGL-20::GFP is represented from high (red) to low (blue). (I) Corrected total EGL-20::GFP fluorescence in each zone of comma stage embryos. EGL-20::GFP levels in the *pdi-1(gk271)* mutant are higher in zone 1 and lower in zones 2 and 3 compared to wild type animals. Data are expressed as mean ± s.d. and statistical significance was assessed by Welch’s t test. n = 15, *p<0.05, **p<0.001.

### PDI-1 and EGL-20/Wnt Function in the Same Genetic Pathway to Regulate Neuronal Migration

We next used genetic analysis to determine whether *pdi-1* and *egl-20* act in the same pathway to direct HSN development (Figure 3F and Table S1). We examined the *egl-20(n585)* missense allele in which a conserved cysteine is substituted to serine (C99S) (Maloof et al., 1999). Mutation of this amino acid in human Wnt3a causes reduction of secretion and activity, presumably through alteration of globular structure (MacDonald et al., 2014). We show that *egl-20(n585)* causes highly penetrant defects (∼75%) in HSN development (Figure 3F), as previously reported (Desai et al., 1988; Forrester et al., 2004; Harris et al., 1996b). We found that the *pdi-1(gk271); egl-20(n585)* compound mutant had no enhancement of penetrance or expressivity of HSN defects compared to the *egl-20(n585)* single mutant (Figure 3F and Table S1), suggesting that these genes act in the same genetic pathway to regulate HSN development. We considered that the penetrance of HSN defects observed in *egl-20(n585)* animals may have reached a ceiling and, as such, no enhancement of phenotype was possible. To examine this possibility, we introduced a LIN-44/Wnt mutation into the *egl-20(n585)* mutant background. We found that the *egl-20(n585); lin-44(n1972)* compound mutant exhibits increased penetrance of HSN defects when compared to the *egl-20(n585)* single mutant (Figure 3F and Table S1), corroborating a previous study (Zinovyeva et al., 2008). Similarly, we found that the *lin-44(n1972)* mutation enhances the *pdi-1(gk271)* HSN developmental defects (Figure 3F and Table S1). Together, these data show that *pdi-1* acts in the same pathway as *egl-20* and in parallel to *lin-44* to control HSN development.

HSN development is controlled by a complex network of Wnt ligands and Frizzled receptors (Pan et al., 2006; Zinovyeva and Forrester, 2005; Zinovyeva et al., 2008). The Frizzled receptors MIG-1 and CFZ-2 act in a partially overlapping manner to coordinate HSN development, where CFZ-2 function is only revealed when MIG-1 is abrogated (Pan et al., 2006; Zinovyeva et al., 2008). Loss of EGL-20 enhances HSN developmental defects in a *mig-1* mutant (Table S1) but not the *cfz-*2 mutant (Zinovyeva and Forrester, 2005), suggesting that EGL-20 can act on both receptors but predominantly on MIG-1. We examined whether PDI-1 exhibits a similar regulatory relationship as EGL-20 with the Frizzled receptors. First, we found that loss of PDI-1 in a *mig-1* mutant causes fully penetrant HSN migration defects (Table S1). In contrast, the penetrance and expressivity of the *pdi-1(gk271)*; *cfz-2(ok1201)* double mutant is no different from the *pdi-1(gk271)* single mutant (Table S1). Together, these data show that PDI-1 acts through the same signaling pathway as EGL-20 to direct HSN migration.

### PDI-1 Controls EGL-20/Wnt Gradient Formation

The formation of an EGL-20 gradient is crucial for correct HSN and Q cell migration in *C. elegans* (Coudreuse et al., 2006; Pan et al., 2006). As PDI-1 controls EGL-20-dependent neuron migration from EGL-20-producing cells (Figure 3E), we hypothesized that loss of PDI-1 function affects the EGL-20 anteroposterior gradient. Three-dimensional reconstructions of EGL-20::GFP localization in comma stage embryos reveal that EGL-20 forms an anteroposterior gradient, as shown previously (Figure 3H and S3) (Coudreuse et al., 2006; Pan et al., 2006). We show that the EGL-20::GFP gradient is perturbed in *pdi-1(gk271)* mutant animals (Figure 3G-I and S3). Loss of PDI-1 elevates EGL-20 localization immediately adjacent to its site of expression with concomitant weaker distal expression (Figure 3G-I and S3). These results reveal that PDI-1 is involved in a key phase of EGL-20 maturation within EGL-20-expressing cells to enable formation of an EGL-20::GFP gradient.

### A PDI Inhibitor Diminishes Wnt3a Secretion from Human Cells

Most mammalian genomes harbor 19 Wnt and over 20 PDI proteins that exhibit diverse expression domains and biological functions (Ellgaard and Ruddock, 2005; Galligan and Petersen, 2012; Willert and Nusse, 2012). Mammalian Wnt3a secretion is abrogated when individual cysteine residues are substituted to alanine (MacDonald et al., 2014), suggesting that Wnt secretion is influenced by the presence of correctly coordinated disulfide bonds. We therefore examined if the role of PDIs in Wnt secretion in *C. elegans* is conserved in human cells by inhibiting PDI function with the 16F16 small-molecule PDI inhibitor (Hoffstrom et al., 2010; Kaplan et al., 2015). Using immunofluorescence, we assayed endogenous Wnt3a levels in human embryonic kidney cells (HEK293T) and found that PDI inhibition caused cellular accumulation of Wnt3a (Figure 4A-B). Elevated Wnt3a protein in HEK293T cells was not due to an increase in expression, as Wnt3a mRNA levels in 16F16-treated cells are unchanged (Figure 4C). To assess Wnt3a secretion, we transfected HEK293T cells with C-terminally V5-tagged Wnt3a (Wnt3a-V5) (Figure 4D-E). We found that the 16F16 PDI inhibitor reduced Wnt3a-V5 secretion in a dose-responsive manner (Figure 4D-E), with no detectable effect on cell viability (Figure S4). To confirm the impact of 16F16 PDI inhibition on secretion is specific to Wnt3a, and not secretion in general, we expressed FLAG-tagged TIG-2 (a bone morphogenetic secreted protein) in HEK293T cells and found that the 16F16 inhibitor had no detectable effect on secretion (Figure S5). These data demonstrate that, as in the *C. elegans* embryo, PDIs are important for correct Wnt secretion from human cells and are therefore fundamental regulators of Wnt signaling.

**Figure 4.**
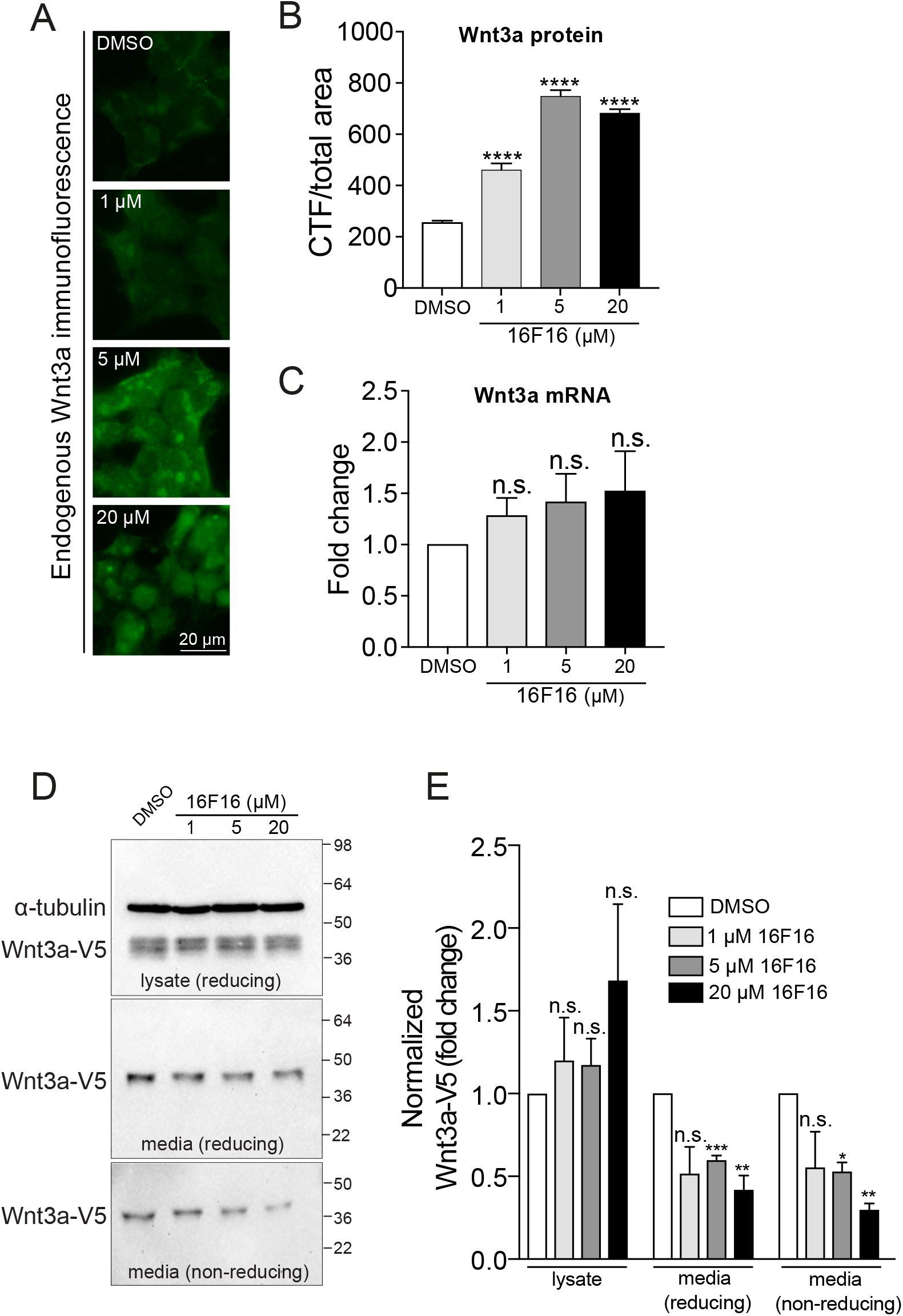
A PDI inhibitor reduces Wnt3a secretion. (A-C) The 16F16 PDI inhibitor increases the level of endogenous Wnt3a protein without affecting transcription. (A) Endogenous Wnt3a immunofluorescence in HEK293T cells incubated with increasing concentrations (0, 1, 5 and 20 µM) of 16F16 in DMSO. Scale bar 20 µm. (B) Quantification of endogenous Wnt3a immunofluorescence from (A) shows increased levels of Wnt3a protein. Data are expressed as mean ± s.d. and statistical significance was assessed by ANOVA followed by Dunnett’s Multiple Comparison Test. n>150 cells from three independent experiments. ****p<0.0001 compared to DMSO control. (C) Wnt3a transcription in HEK293T cells is not affected by incubation with 16F16. Data are expressed as mean ± s.d. and statistical significance was assessed by ANOVA followed by Dunnett’s Multiple Comparison Test. Measurements were taken in triplicate. n.s. – not significant compared to DMSO control. (D-E) Wnt3a-V5 plasmid was transfected into HEK293T cells and then incubated with increasing concentrations (0, 1, 5 and 20 µM) of 16F16 in DMSO. Western blot analysis was performed using anti-V5 (to detect Wnt3a-V5) and anti-a-tubulin (control). Three Western blots from independent samples were performed, a representative example of which is shown in (D), and mean values depicted in (E). Data are presented as DMSO-control set to one-fold. Data are expressed as mean ± s.d. and statistical significance was assessed by Welch’s t test. *p<0.02, **p<0.07, ***p<0.001, n.s. – not significant compared to DMSO-treated cells.

## Discussion

Wnt proteins are lipid-modified morphogens that can form long-range concentration gradients to control developmental patterning. Multiple regulatory mechanisms have evolved to control the amplitude, range and precision of Wnt-directed events. A fundamental question is how Wnt protein folding and maturation is controlled in Wnt-producing cells to enable secretion. Previous studies found that Wnt secretion is controlled by the function of the retromer complex and the Wnt-binding protein WIs (Banziger et al., 2006; Coudreuse et al., 2006; Yang et al., 2008). Here, we report the discovery of a novel regulatory mechanism through which Wnt secretion is controlled by protein disulfide isomerases.

The complexity of Wnt protein secondary structure is coordinated by intracellular disulfide bonds between conserved and invariantly positioned cysteine residues. We reasoned that specific protein chaperones would control disulfide bond formation and that this would be important to control Wnt protein secretion. Through genetic screening of candidate molecules, we identified a protein disulfide isomerase PDI-1 that is crucial for EGL-20/Wnt protein secretion in *C. elegans*. As such, in the absence of PDI-1, EGL-20/Wnt gradient formation is diminished. To examine the phenotypic consequence of a reduced EGL-20/Wnt gradient in *pdi-1* mutant animals, we analyzed two well-characterized EGL-20/Wnt-dependent neurodevelopmental events. We found that PDI-1 controls the migratory capacity of the HSN and Q cells and performs this function cell-non-autonomously from a discrete subset of epidermal cells that co-express EGL-20/Wnt. However, expressing PDI-1 in adjacent cells that express LIN-44/Wnt did not rescue the *pdi-1* mutant phenotype. This suggests a specific function for PDI-1 in EGL-20/Wnt-expressing cells. We further showed that a pharmacological inhibitor of PDIs phenocopies the HSN developmental defects caused by loss of PDI-1 but does not enhance the defects of the *pdi-1(gk271)* null mutant. This strongly suggests that this PDI inhibitor can abrogate PDI-1 function *in vivo* in *C. elegans*. We used the same PDI inhibitor in human cells to show that the mechanism we revealed is conserved. Here, we found that PDI inhibition also decreases Wnt3a secretion from human cells but does not affect release of a BMP morphogen. This may suggest that BMPs are not regulated by PDIs and that disulfide bond formation is not critical for BMP secretion.

A previous study elegantly examined the importance of disulfide bonds for secretion of human Wnt3a from HEK cells (MacDonald et al., 2014). This *in vitro* system showed that substitution of specific, but not all, cysteine residues with alanine reduced Wnt3a secretion (MacDonald et al., 2014). This suggests that the location of unpaired cysteines has differential effects on the ability of Wnt3a to be secreted. Studies also showed that wild-type Wnt3a can form erroneous disulfide bonds during maturation, which would need to be reconfigured prior to secretion (MacDonald et al., 2014; Zhang et al., 2012). Our *in vivo* study has revealed that this function is likely performed by disulfide isomerases.

Sequencing of Wnt mutations isolated in model organism forward genetic screens and in human disease states also reveals the importance of cysteine residues for Wnt protein function. Cysteine mutations in the *Drosophila melanogaster* Wnt protein Wingless, C104S and C242Y, causes temperature-sensitive deficit in secretion and null phenotype, respectively (Dierick and Bejsovec, 1998; Vandenheuvel et al., 1993). For *C. elegans* EGL-20/Wnt, the C99S missense mutation behaves as a null allele whereas the C110Y and C166Y mutations are hypomorphic (Desai et al., 1988; Maloof et al., 1999). In human studies, cysteine mutations in diverse Wnt proteins occur in disease. In Robinow syndrome, which is characterized by short-limbed dwarfism, a causative mutation in Wnt5a (C182R) and other mutations (C69Y and C83S) have been identified (Person et al., 2010; Robinow et al., 1969; Roifman et al., 2015). In addition, a heterozygous Wnt10b mutation (C256Y) was discovered as a potential obesity-related lesion (Christodoulides et al., 2006). Crucially, these mutations have been shown to be important for Wnt secretion and activity *in vitro* (Christodoulides et al., 2006; MacDonald et al., 2014). Together, these studies in animal models and human disease screening support the finding that individual cysteines have differential effects on Wnt function and secretion (MacDonald et al., 2014). The differences observed may be due redundancy in the Wnt system, the specific complement of Wnt proteins that are required to drive certain developmental and homeostatic processes and/or genetic background concerned.

Our genetic data show that loss of PDI-1 in *C. elegans* does not precisely phenocopy the EGL-20/Wnt null (Figure 3). Therefore, a proportion of EGL-20/Wnt protein is likely secreted and functional in *pdi-1* loss-of-function animals. This suggests that additional chaperones are required for folding and secretion of Wnts in laboratory conditions. Further studies are therefore required to identify the molecules concerned.

In conclusion, Wnt protein structure is coordinated by multiple disulfide bonds which have differential importance for Wnt secretory capacity. During maturation, nascent Wnt proteins can form inappropriate disulfide bonds (MacDonald et al., 2014; Zhang et al., 2012) that need to be resolved prior to release to enable appropriate Wnt signaling. It is also possible that variations in disulfide bond arrangement may provide flexibility for interactions between Wnts and specific receptor molecules. Therefore, the oxidation and rearrangement of disulfide bonds is a vital and pervasive step during Wnt biogenesis – a function we have shown to be mediated by protein disulfide isomerases.

## Supplementary Material

### Supplementary Figures

**Figure S1.**
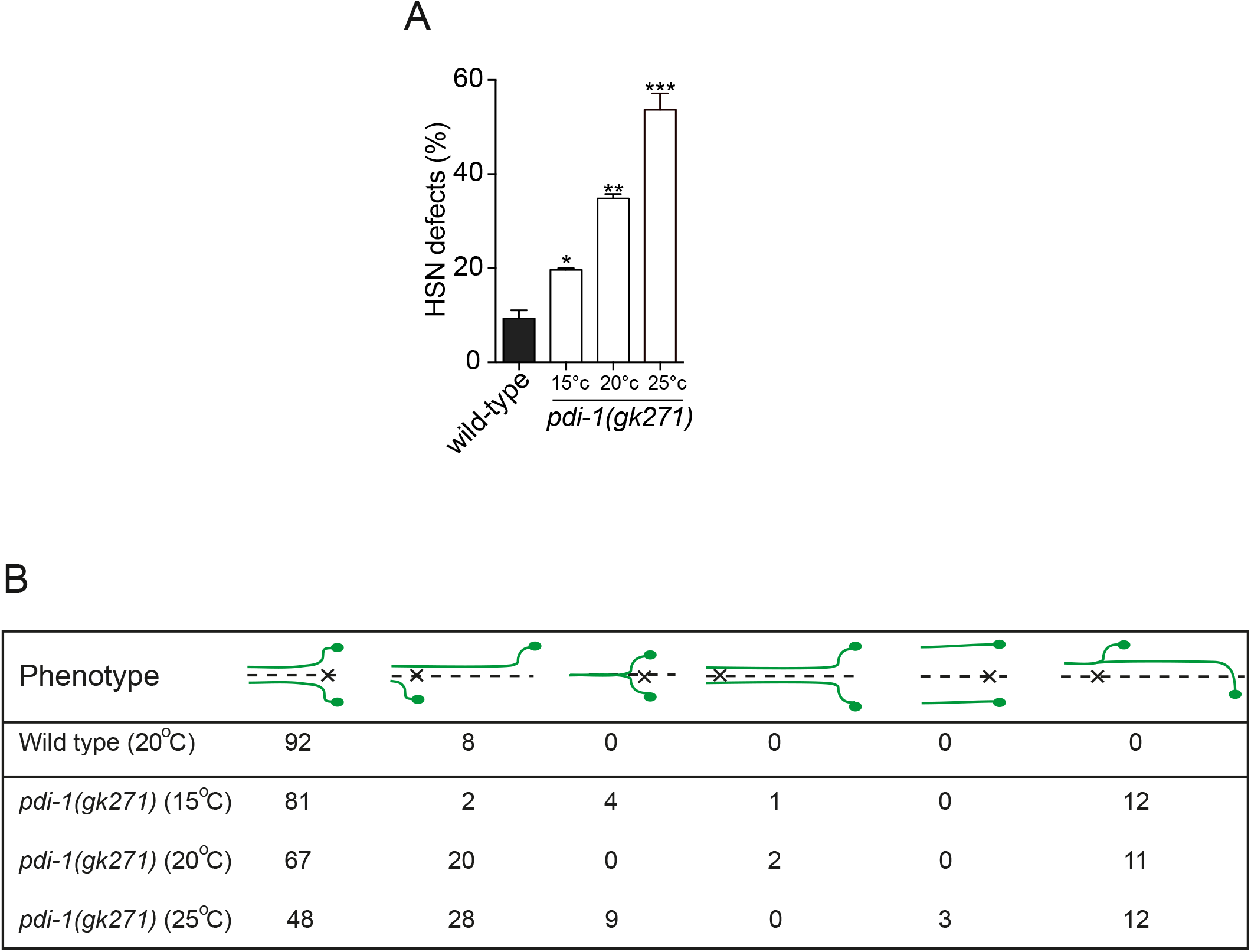
Related to Figure 1. *pdi-1(gk271)* HSN Defects are Temperature-Sensitive. (A) HSN developmental defects of *pdi-1(gk271)* mutant animals are more severe at elevated incubation temperature. Data are expressed as mean ± s.d. and statistical significance was assessed by ANOVA followed by Dunnett’s Multiple Comparison Test. n>100, *p<0.01, **p<0.005, ***p<0.001. *pdi-1(gk271)* mutant animals were incubated at 15°C, 20°C or 25°C. Defects in wild-type animals at 20°C are shown for comparison, which do not significantly alter with incubation at 25°C (13% penetrance, n = 30). (B) Phenotypic breakdown of HSN developmental defects observed in *pdi-1(gk271)* mutant animals incubated at 15°C, 20°C and 25°C. Wild type defects at 20°C are shown for comparison. The percentage of animals exhibiting each phenotype is indicated. Dotted line represents the ventral midline and the ‘X’ represents the vulva. n>100.

**Figure S2.**
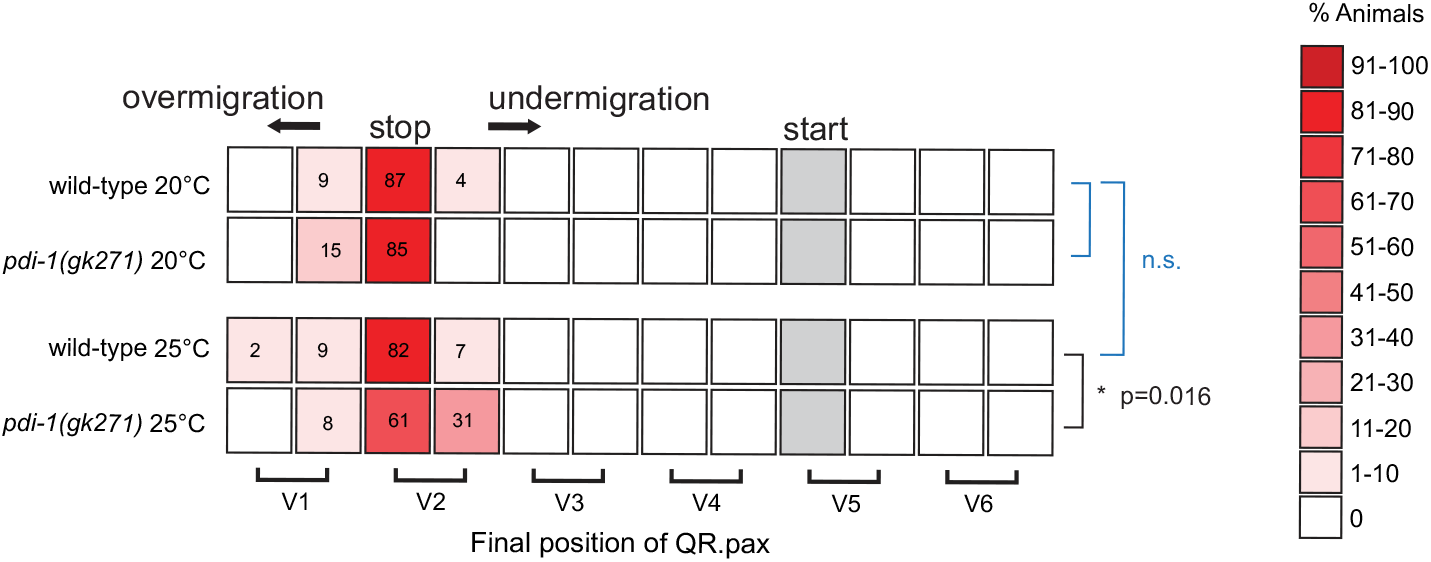
Related to Figure 1. Nomarski Scoring of QR.pax Migration. Average position of the QR.pax neuroblast with respect to seam cells V1.a to V6.p in wild-type and *pdi-1(gk271)* animals cultivated at 20°C and 25°C. Values listed are percentiles of the total number of scored cells; the red coded heat map displays the range of percentile values. Statistical significance was calculated using Fisher’s exact test. n>30 for each cell-type scored, * p<0.016, n.s. – not significant.

**Figure S3.**
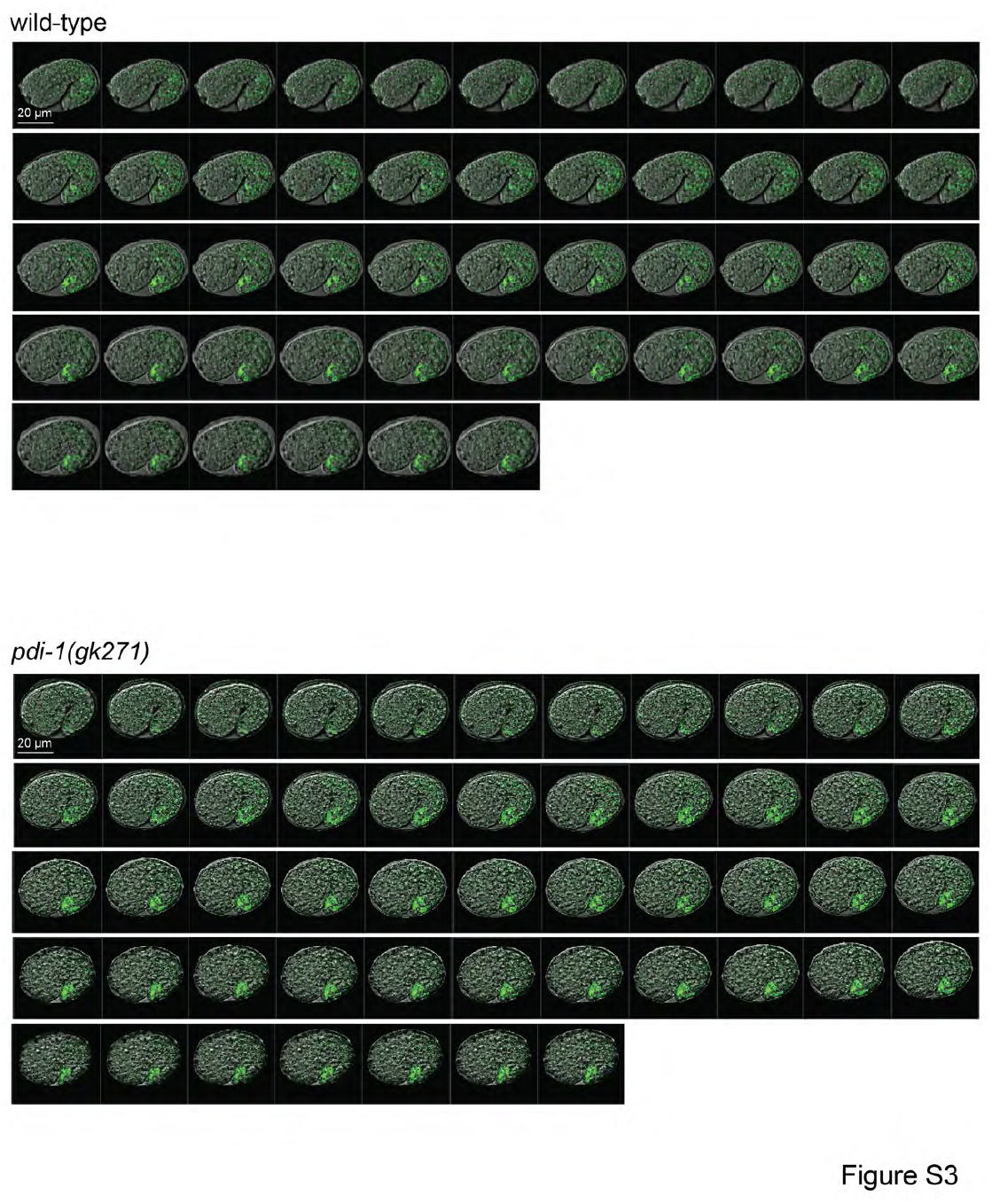
Related to Figure 3. Confocal Analysis of EGL-20::GFP secretion. Confocal images of EGL-20::GFP expression in wild-type and *pdi-1(gk271)* mutant embryos. Images were used to generate 3D-reconstructed heat maps shown in Figure 3H. Each image represents confocal slices that are 0.22µm apart

**Figure S4.**
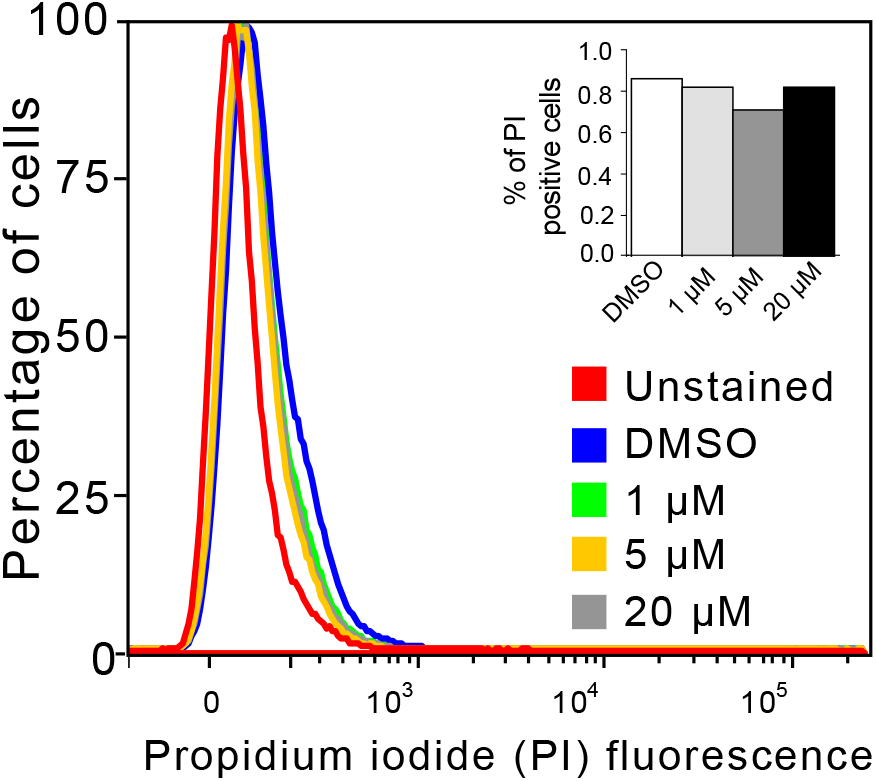
Related to Figure 4. The 16F16 PDI Inhibitor does not Cause HEK293T Cell Death. HEK293T cells treated for 16 h with 16F16 (1, 5 or 20 µM) exhibit no increase in cell death compared to DMSO-treated cells, as measured by propidium iodide (PI) staining. Insert shows the percentage of PI-stained cells in each condition. n>100,000 cells.

**Figure S5.**
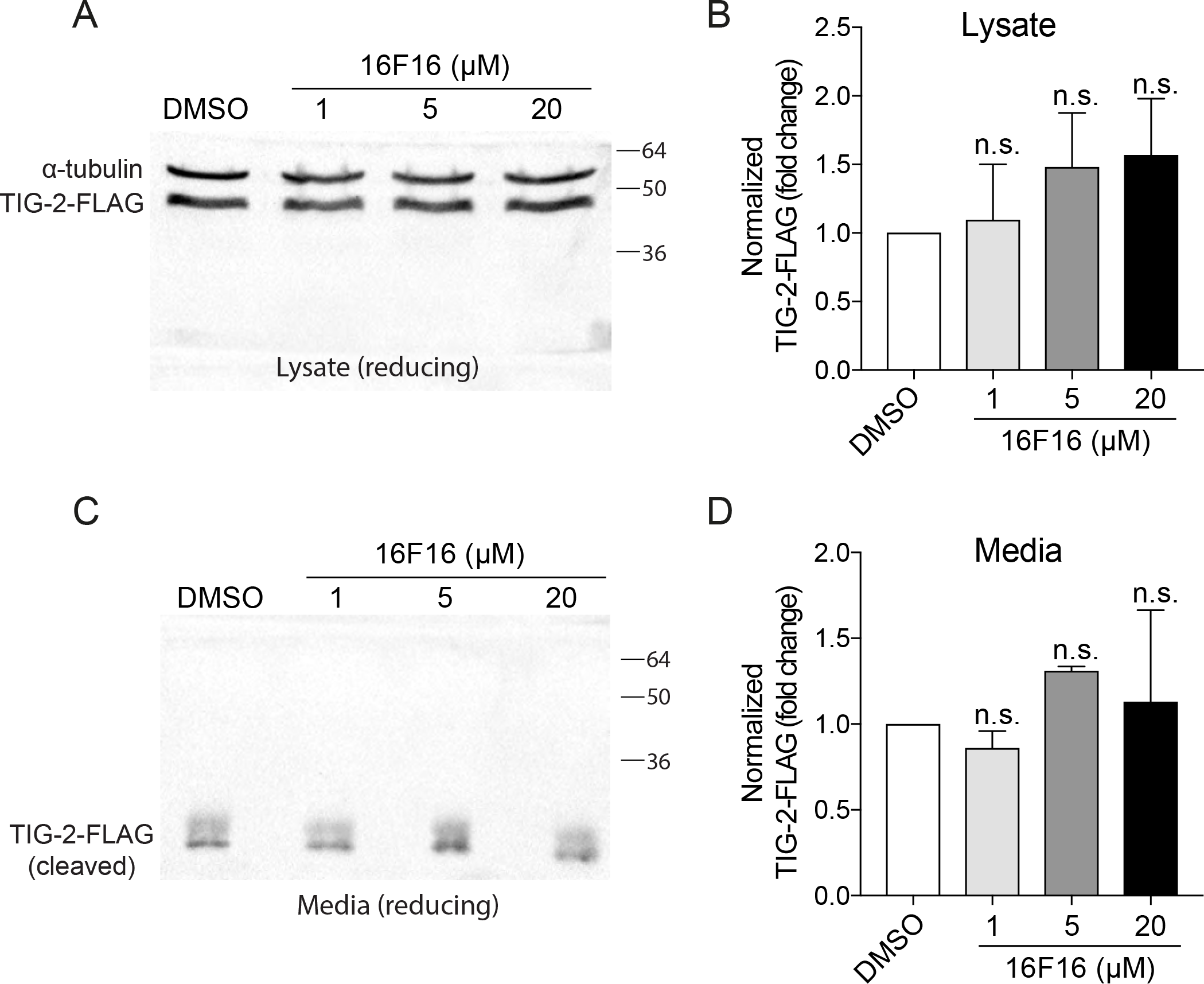
Related to Figure 4. The 16F16 PDI Inhibitor does not Affect TIG-2-FLAG Release from HEK Cells. (A-D) A TIG-2-FLAG plasmid was transfected into HEK293T cells and then incubated with increasing concentrations (0, 1, 5 and 20 µM) of 16F16 in DMSO. Western blot analysis was performed using anti-FLAG (to detect TIG-2-FLAG) and anti-a-tubulin (control). Three Western blots from independent samples were performed, a representative example of which is shown in (A – lysate and C – media), and mean values depicted in (B – lysate and D – media). Data are presented as DMSO-control set to one-fold. Data are expressed as mean ± s.d. and statistical significance was assessed by Welch’s t test. n.s. – not significant compared to DMSO-treated cells. Note that the molecular weight of TIG-2-FLAG in the media is lower than in the lysate as bone morphogenetic proteins, such as TIG-2, undergo proteolytic cleavage within the secretory pathway prior to release from cells.

### Tables

**Table S1.**
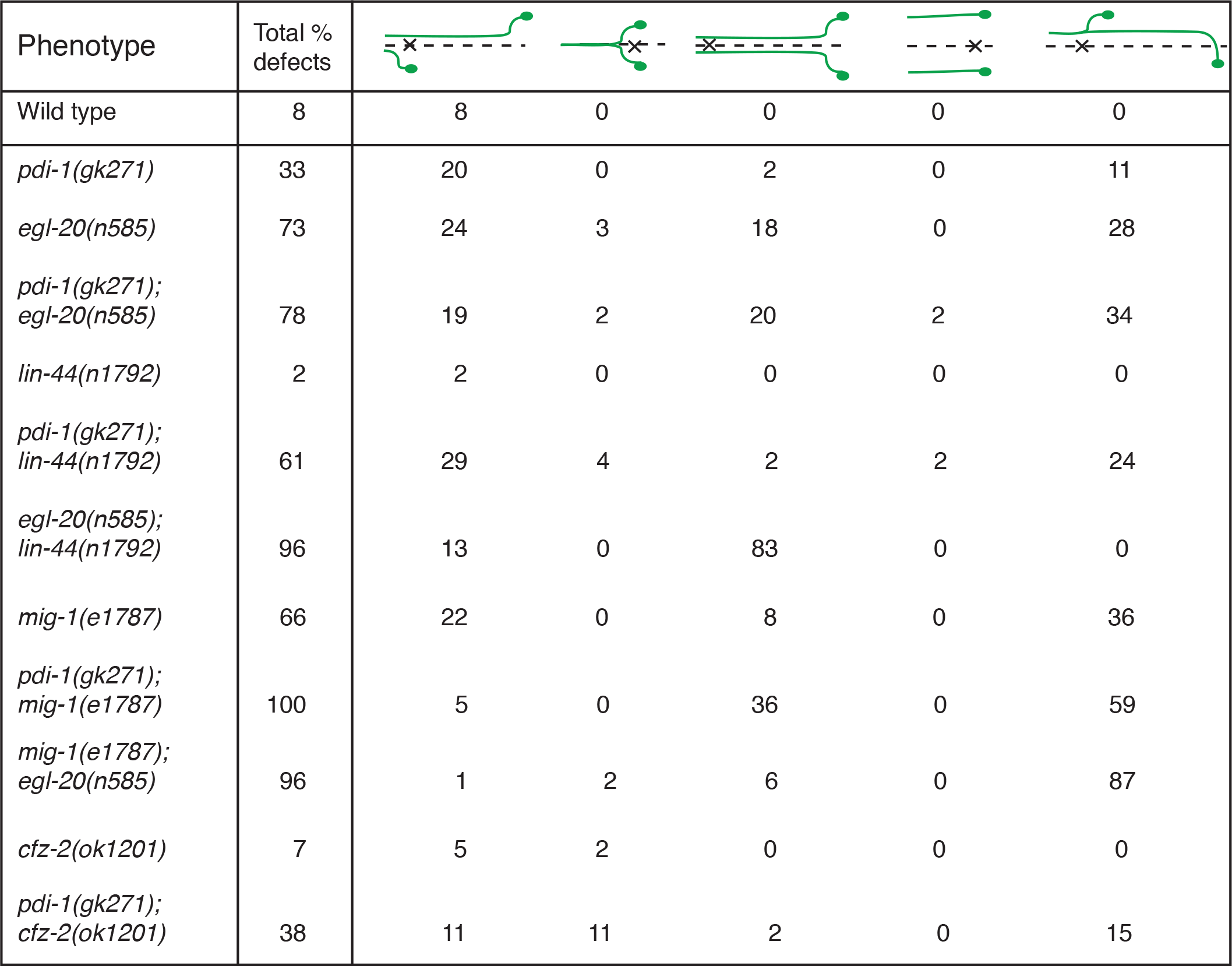
Double mutant analysis. Phenotypic breakdown of HSN developmental defects observed in single and double mutant combinations of animals lacking PDI-1, Wnt morphogens (EGL-20 or LIN-44) or Frizzled receptors (MIG-1 or CFZ-2). These data show that PDI-1 acts in the same pathway as EGL-20 and CFZ-2 and in parallel to LIN-44 and MIG-1. The percentage of animals exhibiting each phenotype is indicated. Dotted line represents the ventral midline and the ‘X’ represents the vulva. n>100.

| Phenotype | Total % defects |  |  |  |  |  |
| --- | --- | --- | --- | --- | --- | --- |
| Wild type | 8 | 8 | 0 | 0 | 0 | 0 |
| *pdi-1(gk271)* | 33 | 20 | 0 | 2 | 0 | 11 |
| *egl-20(n585)* | 73 | 24 | 3 | 18 | 0 | 28 |
| *pdi-1(gk271); egl-20(n585)* | 78 | 19 | 2 | 20 | 2 | 34 |
| *lin-44(n1792)* | 2 | 2 | 0 | 0 | 0 | 0 |
| *pdi-1(gk271); lin-44(n1792)* | 61 | 29 | 4 | 2 | 2 | 24 |
| *egl-20(n585); lin-44(n1792)* | 96 | 13 | 0 | 83 | 0 | 0 |
| *mig-1(e1787)* | 66 | 22 | 0 | 8 | 0 | 36 |
| *pdi-1(gk271); mig-1(e1787)* | 100 | 5 | 0 | 36 | 0 | 59 |
| *mig-1(e1787); egl-20(n585)* | 96 | 1 | 2 | 6 | 0 | 87 |
| *cfz-2(ok1201)* | 7 | 5 | 2 | 0 | 0 | 0 |
| *pdi-1(gk271); cfz-2(ok1201)* | 38 | 11 | 11 | 2 | 0 | 15 |

**Table S2.**
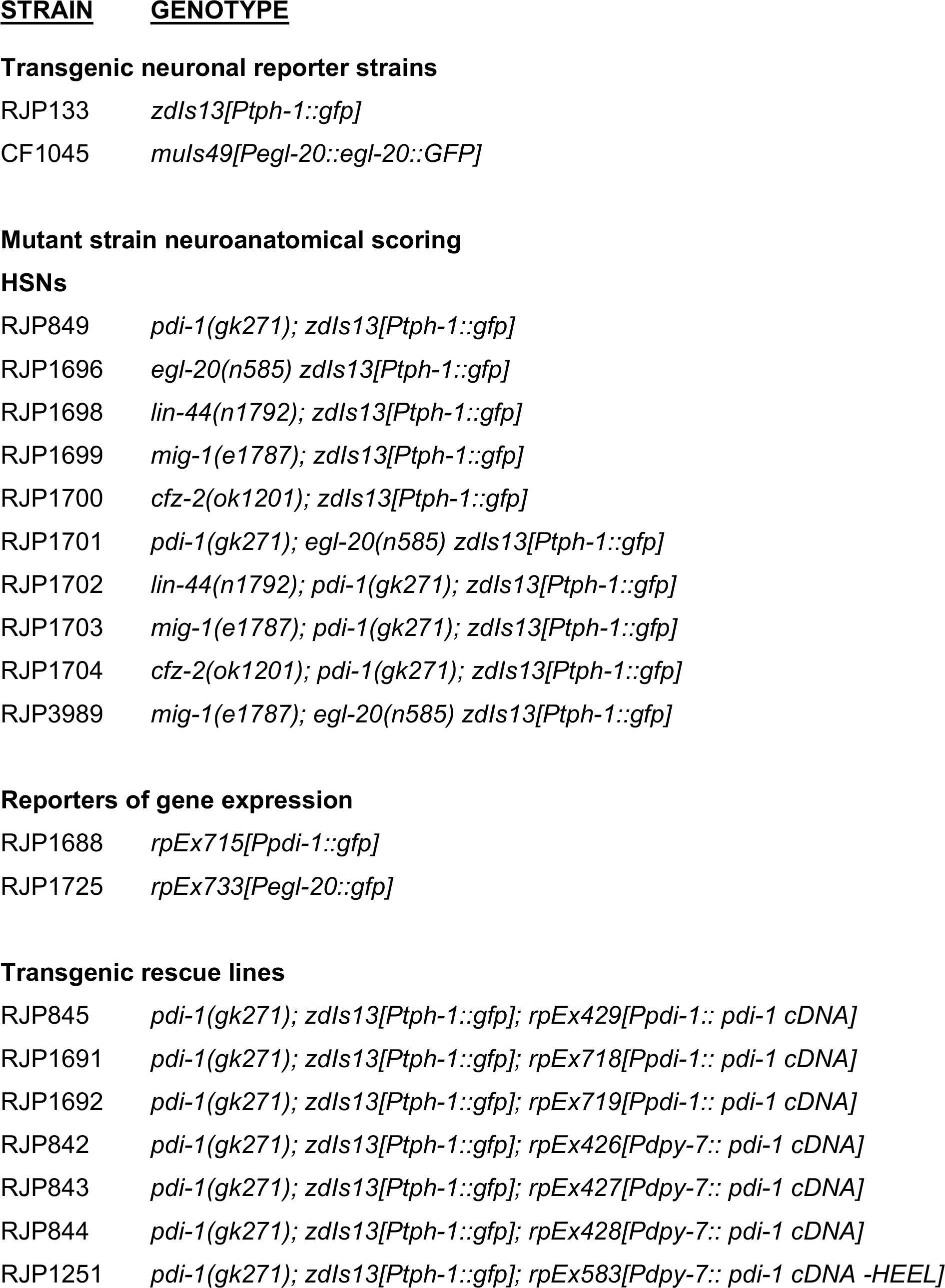
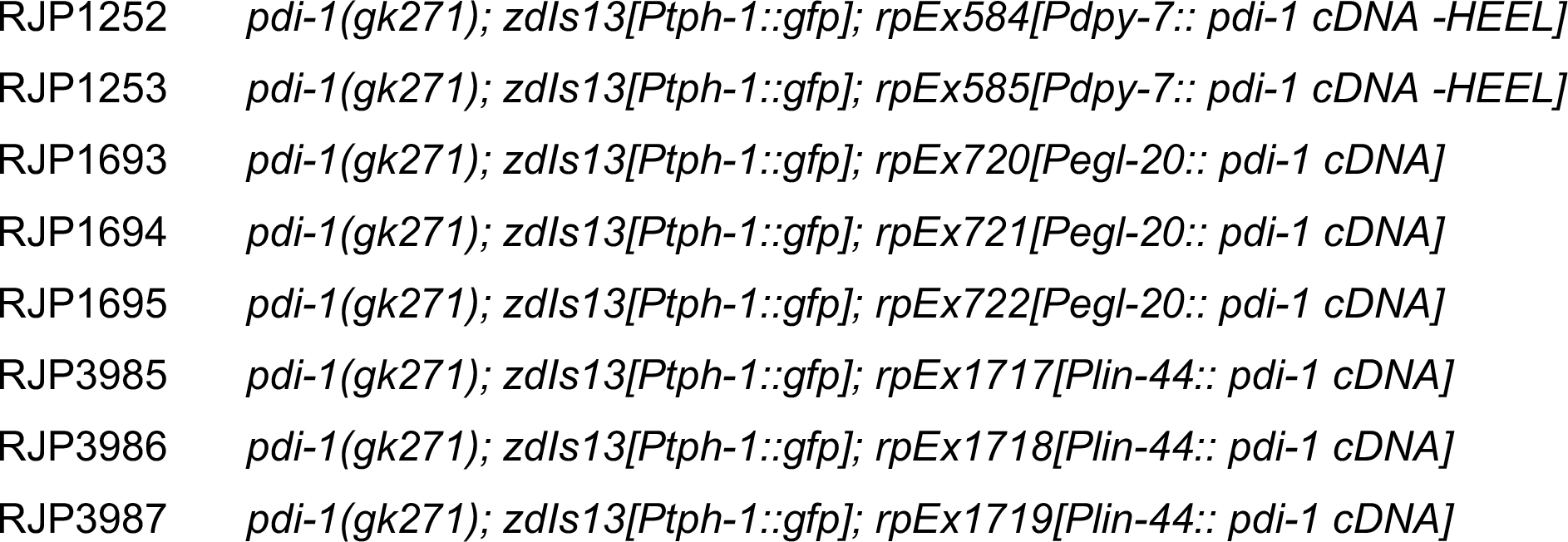
Strains used in this study.

**Table S3.**
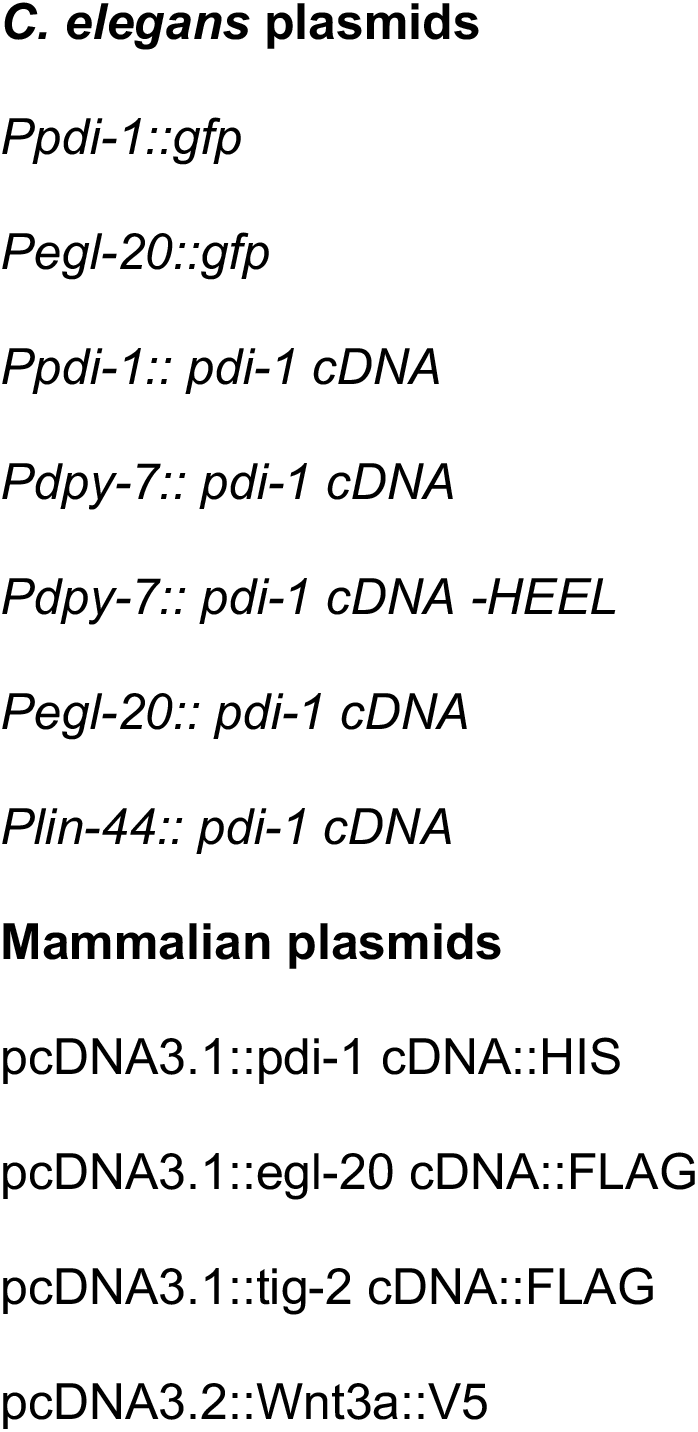
Plasmids used in this study.

## Experimental Procedures

### Mutant and transgenic reporter strains

Strains were grown using standard growth conditions on NGM agar at 20^0^C on *Escherichia coli* OP50, unless otherwise stated (Brenner, 1974).

Neuroanatomical reporter strains used – *mgIs70 Is[Ptph-1::gfp]*; LGIV *zdIs13*: *Is[Ptph-1::gfp]*. Detailed strain information is detailed in Table S2.

### Molecular cloning

#### *Ppdi-1::pdi-1* cDNA rescue construct

The *Ppdi-1::pdi-1* cDNA rescue construct was generated by cloning the 841 bp *pdi-1* promoter with SphI-SmaI into the pPD49.26 vector and then the *pdi-*1 cDNA using NheI-SacI.

#### *Pdpy-7::pdi-1* cDNA rescue construct

The *Pdpy-7::pdi-1* cDNA rescue construct was generated by cloning *pdi-*1 cDNA with NheISacI into a *dpy-7* expression vector.

#### *Pdpy-7::pdi-1* cDNA – HEEL rescue construct

The *Pdpy-7::pdi-1* cDNA –HEEL rescue construct was generated using site-directed mutagenesis to remove the 12 nucleotides that encode HEEL at the C-terminus of the PDI-1 protein.

#### *Pegl-20::pdi-1* cDNA rescue construct

The *Pegl-20::pdi-1* cDNA rescue construct was generated by cloning the 1910 bp *egl-20* promoter with SphI-SmaI into the pPD49.26 vector containing the *pdi-*1 cDNA.

#### *Plin-44::pdi-1* cDNA rescue construct

The *Plin-44::pdi-1* cDNA rescue construct was generated by cloning the 1150 bp *lin-44* promoter with SphI-SmaI into the pPD49.26 vector containing the *pdi-*1 cDNA.

#### *egl-20::gfp* reporter construct

The *Pegl-20::gfp* reporter construct was generated by cloning the 1910 bp *egl-20* promoter with SphI-SmaI into the promoterless *gfp* pPD95.75 expression vector.

#### pcDNA3.1(zeo)::pdi-1 cDNA::HIS tag

The *pdi-1*cDNA::HIS mammalian expression construct was generated by cloning the 1458 bp *pdi-1* cDNA with NheI-KpnI into the pcDNA3.1(zeo) expression vector.

#### pcDNA3.1(zeo)::egl-20 cDNA::FLAG tag

The *egl-20*cDNA::FLAG mammalian expression construct was generated by cloning the 1182 bp *egl-20* cDNA with NheI-KpnI into the pcDNA3.1(zeo) expression vector.

#### pcDNA3.1(zeo)::tig-2 cDNA::FLAG tag

The *tig-2*cDNA::FLAG mammalian expression construct was generated by cloning the 1098 bp *tig-2* cDNA with NheI-HindIII into the pcDNA3.1(zeo) expression vector.

#### DNA constructs and transgenic lines

Rescue constructs were injected into the *pdi-1(gk271)* mutant background at 5–15 ng/µl with *Pmyo-2::mcherry* (5 ng/µl) as injection marker. Expression constructs were injected into N2 background at 50 ng/µl with *Pmyo-2::mcherry* (5 ng/µl) as injection marker. Microinjections were performed using standard methods (Mello et al., 1991).

#### RNAi experiments

RNAi feeding experiments were conducted following the Ahringer lab protocol (Fraser et al., 2000). L4440 vectors (with or without specific dsRNA) were amplified in *Escherichia coli* HT115 and selected with ampicillin and tetracycline. 3–5 L4 hermaphrodites were moved into each RNAi bacteria seeded plate and progeny were scored after 4 days as young adults. For each knockdown strain, three different experiments were conducted on three independent days.

#### PDI inhibition in *C. elegans*

A 4 µM solution of the 16F16 small-molecule PDI inhibitor (Sigma – SML0021) was made using M9 buffer (0.1% DMSO final concentration). Synchronized L4 hermaphrodites were incubated in 10 ml of 16F16 (4 µM) in M9 supplemented with 300 µl of concentrated OP50 *E. coli*. For the control experiment, worms were incubated in 10 ml of M9 (0.1% DMSO) plus 300 µl of concentrated OP50 *E. coli*. Worms were shaken at 20°C for 48 hours, after which they were centrifuged at 2500 rpm for 1 minute and then transferred to standard NGM plates seeded with OP50 *E. coli*. HSN anatomy was scored the following day.

#### Fluorescence microscopy

Animals were anesthetized with 20 mM NaN_3_ on 5% agarose pads, and images were obtained with an Axio Imager M2 fluorescence microscope and Zen software (Zeiss).

#### Nomarski analysis of long-range migrating neurons

The final position of Q cell descendants (QR.pax and QL.pax) and HSNs were determined using Nomarski optics in late L1 larvae, as described (Coudreuse et al., 2006). The positions of QR.pax, QL.pax and HSNs were determined with respect to the seam cell daughters V1.a to V6.p.

#### 3D analysis of EGL-20::GFP distribution in *C. elegans* embryos

Comma stage wild-type and *pdi-1(gk271)* embryos expressing EGL-20::GFP were imaged with a Leica SP5 Confocal microscope using a HC PL APO 63x/1.4 oil objective. We imaged slices of 0.22 µm that were processed using Adobe illustrator CC 2018. 3D models of embryos were generated using Imaris 9.1.2 3D interactive microscopy analysis software from Bitplane (Gopal et al., 2017). The intensity of EGL-20::GFP was calculated after projecting the 3D slices to a 2D plane using FIJI image analysis software (Figure S3). To score the distribution of EGL-20::GFP, embryos were divided into three zones. Zone 1 encompasses the immediate vicinity of EGL-20::GFP producing cells, with Zones 2 and 3 covering more distal regions. Background values were obtained from an area of the embryo without detectable EGL-20::GFP.

#### Endogenous Wnt3a staining of HEK293T cells

HEK293T cells were cultured overnight in 12-well tissue culture plates in DMEM containing 10% serum and 2.5 mM L-glutamine before treating with 250 µl media containing 16F16 or DMSO for 16 h. Cells were fixed using 4% paraformaldehyde and stained with anti-Wnt3a antibody and imaged using Zeiss PLAN-APOCHROMAT 40×/1.4 objective. Wnt3a immunofluorescence was quantified using FIJI with the following formula: Corrected total fluorescence (CTF) = Integrated Density – (Area of selected cell x Mean fluorescence of background readings). Wnt3a level from at least 150 cells from three independent experiments were measured.

#### Wnt3a secretion assay in HEK293T cells

HEK293T cells were cultured at 37°C in 12-well tissue culture plates in DMEM containing 10% serum and 2.5 mM L-glutamine. Cells were transfected with Wnt3aV5 or TIG-2-FLAG using Lipofectamine 2000 (Thermo Fisher, 11668019) according to manufactures instructions. After 24 h, the media was replaced with 250 µl of serum-free DMEM containing the 16F16 PDI inhibitor (Sigma, SML0021) or DMSO. Cells were incubated at 37°C for 16 h before cells and media were collected. The cells were lysed with 50 µl of 2.5x LDS sample buffer (Thermo Scientific, B0007) containing reducing agent (Thermo Scientific, NP0009). 20 µl of 2.5× LDS sample buffer, with or without reducing agent, was added to the collected media. Samples were run on an SDS-PAGE gel before blotting to a PVDF membrane. Membranes were probed with antibodies against FLAG (Sigma, F1804) and V5-tag (Bio-Rad, MCA1396GA), and a-tubulin (Developmental Studies Hybridoma Bank, 12G10 anti-a-tubulin) before applying HRP conjugated secondary antibodies (Thermoscientific). Blots were developed using BioRad ChemiDoc XRS+. Wnt3a bands were quantified using FIJI image analysis software and values were normalized first against a-tubulin and then against DMSO-treated samples.

#### Fluorescence-activated cell sorting

Cell death after 16F16 treatment of HEK293T was analyzed by propidium iodide (PI) staining followed by FACS. Cells were cultured in 10 cm dishes and were treated with 16F16 (1, 5 or 20 µM) or DMSO for 16 h, after which cells were harvested using dissociation buffer and stained with PI at a ratio of 1:50. The percentage of PI-stained cells was determined using FACS. PI was maintained in the solution during the analysis.

#### qPCR assays

Total RNA was isolated using using the RNAeasy mini kit (Qiagen 74104), according to manufacturer’s instructions. Total cDNA was obtained using oligodT primers and the ImProm-II™ Reverse Transcription System (A3800) followed by quantitative PCR using SYBR green (Thermo Scientific 4385610) and Light Cycler 480 (Roche). The human Ets2 reference gene was used as a control.

#### Primary antibodies

Monoclonal antibodies used in this study were mouse anti-a-tubulin (clone 12G10; DSHB) and mouse anti-V5 (clone SV5-pk1; BioRad MCA1630). The polyclonal antibody used was rabbit anti-Wnt3a (Abcam ab199925).

#### Statistical analysis

Statistical analysis was performed in GraphPad Prism 7 using one-way analysis of variance (ANOVA) for comparison followed by Dunnett’s Multiple Comparison Test or Tukey’s Multiple Comparison Test, where applicable. Values are expressed as mean ± s.d. Differences with a *P* value <0.05 were considered significant. Statistical analysis of HSN and QR/QL descendants was performed using Fisher’s exact test. A Monte Carlo approximation, iterated 10.000 times using SPSStatistics version 22, was used to evaluate significance. Statistical analysis of EGL-20::GFP levels and Wnt-3a secretion was performed using Welch’s t test.

## Acknowledgements

We thank members of the Pocock Laboratory and Brent Neumann for comments on the manuscript. Some strains were provided by the *Caenorhabditis* Genetics Center (University of Minnesota), which is funded by NIH Office of Research Infrastructure Programs (P40 OD010440). This work was supported by the following grants: Monash University Senior Postdoctoral Fellowship to S.G., NWO-ALW (Open Program Grant 822.02.012) to H.C.K., European Research Council (ERC Starting Grant 260807 to R.P.), Lundbeck Foundation (Project R67-A6094 to R.P), NHMRC (Project GNT1105374 to R.P.) and veski Innovation Fellowship (VIF23 to R.P.).

## References

Banziger, C., Soldini, D., Schutt, C., Zipperlen, P., Hausmann, G., and Basler, K. (2006). Wntless, a conserved membrane protein dedicated to the secretion of Wnt proteins from signaling cells. Cell 125, 509–522.

Barrott, J.J., Cash, G.M., Smith, A.P., Barrow, J.R., and Murtaugh, L.C. (2011). Deletion of mouse Porcn blocks Wnt ligand secretion and reveals an ectodermal etiology of human focal dermal hypoplasia/Goltz syndrome. P Natl Acad Sci USA 108, 12752–12757.

Belenkaya, T.Y., Wu, Y.H., Tang, X.F., Zhou, B., Cheng, L.Q., Sharma, Y.V., Yan, D., Selva, E.M., and Lin, X.H. (2008). The retromer complex influences Wnt secretion by recycling Wntless from endosomes to the trans-Golgi network. Developmental Cell 14, 120–131.

Biechele, S., Cox, B.J., and Rossant, J. (2011). Porcupine homolog is required for canonical Wnt signaling and gastrulation in mouse embryos. Developmental Biology 355, 275–285.

Bocchi, R., Egervari, K., Carol-Perdiguer, L., Viale, B., Quairiaux, C., De Roo, M., Boitard, M., Oskouie, S., Salmon, P., and Kiss, J.Z. (2017). Perturbed Wnt signaling leads to neuronal migration delay, altered interhemispheric connections and impaired social behavior. Nat Commun 8.

Braakman, I., and Bulleid, N.J. (2011). Protein folding and modification in the mammalian endoplasmic reticulum. Annual review of biochemistry 80, 71–99.

Brenner, S. (1974). The genetics of Caenorhabditis elegans. Genetics 77, 71–94.

Cadigan, K.M., Fish, M.P., Rulifson, E.J., and Nusse, R. (1998). Wingless repression of Drosophila frizzled 2 expression shapes the Wingless morphogen gradient in the wing. Cell 93, 767–777.

Caramelo, J.J., and Parodi, A.J. (2007). How sugars convey information on protein conformation in the endoplasmic reticulum. Seminars in cell & developmental biology 18, 732–742.

Christodoulides, C., Scarda, A., Granzotto, M., Milan, G., Dalla Nora, E., Keogh, J., De Pergola, G., Stirling, H., Pannacciulli, N., Sethi, J.K., et al. (2006). WNT10B mutations in human obesity. Diabetologia 49, 678–684.

Clevers, H. (2006). Wnt/beta-catenin signaling in development and disease. Cell 127, 469–480.

Coudreuse, D.Y., Roel, G., Betist, M.C., Destree, O., and Korswagen, H.C. (2006). Wnt gradient formation requires retromer function in Wnt-producing cells. Science 312, 921–924.

Desai, C., Garriga, G., McIntire, S.L., and Horvitz, H.R. (1988). A genetic pathway for the development of the Caenorhabditis elegans HSN motor neurons. Nature 336, 638–646.

Dierick, H.A., and Bejsovec, A. (1998). Functional analysis of Wingless reveals a link between intercellular ligand transport and dorsal-cell-specific signaling. Development 125, 4729–4738.

Ellgaard, L., and Ruddock, L.W. (2005). The human protein disulphide isomerase family: substrate interactions and functional properties. EMBO Rep 6, 28–32.

Fass, D. (2012). Disulfide bonding in protein biophysics. Annual review of biophysics 41, 63–79.

Forrester, W.C., Kim, C., and Garriga, G. (2004). The Caenorhabditis elegans Ror RTK CAM-1 inhibits EGL-20/Wnt signaling in cell migration. Genetics 168, 1951–1962.

Fraser, A.G., Kamath, R.S., Zipperlen, P., Martinez-Campos, M., Sohrmann, M., and Ahringer, J. (2000). Functional genomic analysis of C. elegans chromosome I by systematic RNA interference. Nature 408, 325–330.

Galligan, J.J., and Petersen, D.R. (2012). The human protein disulfide isomerase gene family. Hum Genomics 6, 6.

Gopal, S., Boag, P., and Pocock, R. (2017). Automated three-dimensional reconstruction of the Caenorhabditis elegans germline. Dev Biol.

Grzeschik, K.H., Bornholdt, D., Oeffner, F., Konig, A., Boente, M.D., Enders, H., Fritz, B., Hertl, M., Grasshoff, U., Hofling, K., et al. (2007). Deficiency of PORCN, a regulator of Wnt signaling, is associated with focal dermal hypoplasia. Nat Genet 39, 833–835.

Harris, J., Honigberg, L., Robinson, N., and Kenyon, C. (1996a). Neuronal cell migration in C. elegans: regulation of Hox gene expression and cell position. Development 122, 3117–3131.

Harris, J., Honigberg, L., Robinson, N., and Kenyon, C. (1996b). Neuronal cell migration in C. elegans: regulation of Hox gene expression and cell position. Development 122, 3117–3131.

Hilliard, M.A., and Bargmann, C.I. (2006). Wnt signals and frizzled activity orient anterior-posterior axon outgrowth in C. elegans. Dev Cell 10, 379–390.

Hoffstrom, B.G., Kaplan, A., Letso, R., Schmid, R.S., Turmel, G.J., Lo, D.C., and Stockwell, B.R. (2010). Inhibitors of protein disulfide isomerase suppress apoptosis induced by misfolded proteins. Nat Chem Biol 6, 900–906.

Janda, C.Y., Waghray, D., Levin, A.M., Thomas, C., and Garcia, K.C. (2012). Structural basis of Wnt recognition by Frizzled. Science 337, 59–64.

Kadowaki, T., Wilder, E., Klingensmith, J., Zachary, K., and Perrimon, N. (1996). The segment polarity gene porcupine encodes a putative multitransmembrane protein involved in Wingless processing. Gene Dev 10, 3116–3128.

Kaplan, A., Gaschler, M.M., Dunn, D.E., Colligan, R., Brown, L.M., Palmer, A.G., 3rd, Lo, D.C., and Stockwell, B.R. (2015). Small molecule-induced oxidation of protein disulfide isomerase is neuroprotective. Proc Natl Acad Sci U S A 112, E2245–2252.

Kennedy, L.M., Pham, S.C., and Grishok, A. (2013). Nonautonomous regulation of neuronal migration by insulin signaling, DAF-16/FOXO, and PAK-1. Cell reports 4, 996–1009.

Klassen, M.P., and Shen, K. (2007). Wnt signaling positions neuromuscular connectivity by inhibiting synapse formation in C. elegans. Cell 130, 704–716.

MacDonald, B.T., Hien, A., Zhang, X., Iranloye, O., Virshup, D.M., Waterman, M.L., and He, X. (2014). Disulfide bond requirements for active Wnt ligands. J Biol Chem 289, 18122–18136.

Maloof, J.N., Whangbo, J., Harris, J.M., Jongeward, G.D., and Kenyon, C. (1999). A Wnt signaling pathway controls hox gene expression and neuroblast migration in C. elegans. Development 126, 37–49.

Mello, C.C., Kramer, J.M., Stinchcomb, D., and Ambros, V. (1991). Efficient gene transfer in C.elegans: extrachromosomal maintenance and integration of transforming sequences. Embo J 10, 3959–3970.

Mentink, R.A., Middelkoop, T.C., Rella, L., Ji, N., Tang, C.Y., Betist, M.C., van Oudenaarden, A., and Korswagen, H.C. (2014). Cell intrinsic modulation of Wnt signaling controls neuroblast migration in C. elegans. Dev Cell 31, 188–201.

Miller, J.R. (2002). The Wnts. Genome Biol 3, REVIEWS3001.

Modzelewska, K., Lauritzen, A., Hasenoeder, S., Brown, L., Georgiou, J., and Moghal, N. (2013). Neurons Refine the Caenorhabditis elegans Body Plan by Directing Axial Patterning by Wnts. Plos Biol 11.

Mogk, A., Mayer, M.P., and Deuerling, E. (2002). Mechanisms of protein folding: Molecular chaperones and their application in biotechnology. Chembiochem 3, 807–+.

Pan, C.L., Howell, J.E., Clark, S.G., Hilliard, M., Cordes, S., Bargmann, C.I., and Garriga, G. (2006). Multiple Wnts and frizzled receptors regulate anteriorly directed cell and growth cone migrations in Caenorhabditis elegans. Dev Cell 10, 367–377.

Pedersen, M.E., Snieckute, G., Kagias, K., Nehammer, C., Multhaupt, H.A., Couchman, J.R., and Pocock, R. (2013). An epidermal microRNA regulates neuronal migration through control of the cellular glycosylation state. Science 341, 1404–1408.

Person, A.D., Beiraghi, S., Sieben, C.M., Hermanson, S., Neumann, A.N., Robu, M.E., Schleiffarth, J.R., Billington, C.J., van Bokhoven, H., Hoogeboom, J.M., et al. (2010). WNT5A Mutations in Patients With Autosomal Dominant Robinow Syndrome. Dev Dynam 239, 327–337.

Port, F., Kuster, M., Herr, P., Furger, E., Banziger, C., Hausmann, G., and Basler, K. (2008). Wingless secretion promotes and requires retromer-dependent cycling of Wntless. Nat Cell Biol 10, 178–U148.

Prasad, B.C., and Clark, S.G. (2006). Wnt signaling establishes anteroposterior neuronal polarity and requires retromer in C. elegans. Development 133, 1757–1766.

Robinow, M., Silverman, F.N., and Smith, H.D. (1969). A Newly Recognized Dwarfing Syndrome. Am J Dis Child 117, 645–+.

Roifman, M., Marcelis, C.L., Paton, T., Marshall, C., Silver, R., Lohr, J.L., Yntema, H.G., Venselaar, H., Kayserili, H., van Bon, B., et al. (2015). De novo WNT5A-associated autosomal dominant Robinow syndrome suggests specificity of genotype and phenotype. Clin Genet 87, 34–41.

Sulston, J.E., Schierenberg, E., White, J.G., and Thomson, J.N. (1983). The embryonic cell lineage of the nematode Caenorhabditis elegans. Dev Biol 100, 64–119.

Takada, R., Satomi, Y., Kurata, T., Ueno, N., Norioka, S., Kondoh, H., Takao, T., and Takada, S. (2006). Monounsaturated fatty acid modification of Wnt protein: its role in Wnt secretion. Dev Cell 11, 791–801.

Vandenheuvel, M., Harrymansamos, C., Klingensmith, J., Perrimon, N., and Nusse, R. (1993). Mutations in the Segment Polarity Genes Wingless and Porcupine Impair Secretion of the Wingless Protein. Embo J 12, 5293–5302.

Wang, X.L., Sutton, V.R., Peraza-Llanes, J.O., Yu, Z.Y., Rosetta, R., Kou, Y.C., Eble, T.N., Patel, A., Thaller, C., Fang, P., et al. (2007). Mutations in X-linked PORCN, a putative regulator of Wnt signaling, cause focal dermal hypoplasia. Nat Genet 39, 836–838.

Whangbo, J., and Kenyon, C. (1999). A Wnt signaling system that specifies two patterns of cell migration in C. elegans. Mol Cell 4, 851–858.

Wilkinson, B., and Gilbert, H.F. (2004). Protein disulfide isomerase. Biochim Biophys Acta 1699, 35–44.

Willert, K., Brown, J.D., Danenberg, E., Duncan, A.W., Weissman, I.L., Reya, T., Yates, J.R., 3rd, and Nusse, R. (2003). Wnt proteins are lipid-modified and can act as stem cell growth factors. Nature 423, 448–452.

Willert, K., and Nusse, R. (2012). Wnt proteins. Cold Spring Harb Perspect Biol 4, a007864.

Yang, P.T., Lorenowicz, M.J., Silhankova, M., Coudreuse, D.Y., Betist, M.C., and Korswagen, H.C. (2008). Wnt signaling requires retromer-dependent recycling of MIG-14/Wntless in Wnt-producing cells. Dev Cell 14, 140–147.

Yoshikawa, S., McKinnon, R.D., Kokel, M., and Thomas, J.B. (2003). Wnt-mediated axon guidance via the Drosophila derailed receptor. Nature 422, 583–588.

Zhang, X., Abreu, J.G., Yokota, C., MacDonald, B.T., Singh, S., Coburn, K.L., Cheong, S.M., Zhang, M.M., Ye, Q.Z., Hang, H.C., et al. (2012). Tiki1 is required for head formation via Wnt cleavage-oxidation and inactivation. Cell 149, 1565–1577.

Zinovyeva, A.Y., and Forrester, W.C. (2005). The C. elegans Frizzled CFZ-2 is required for cell migration and interacts with multiple Wnt signaling pathways. Dev Biol 285, 447–461.

Zinovyeva, A.Y., Yamamoto, Y., Sawa, H., and Forrester, W.C. (2008). Complex network of Wnt signaling regulates neuronal migrations during Caenorhabditis elegans development. Genetics 179, 1357–1371.

